# High-density environmental DNA metabarcoding reveals fine-scale three-dimensional structure of a fish community in a semi-closed bay

**DOI:** 10.64898/2026.11.04.764206

**Authors:** Hiroaki Murakami, Ryosuke Nakadai, Toshiaki S. Jo, Qianqian Wu, Daiki Kadotani Tomojiri, Reiji Masuda

## Abstract

Coastal fish assemblages are influenced by environmental heterogeneity across horizontal and vertical dimensions, yet their fine-scale three-dimensional structure remains poorly understood. We investigated spatial variation in fish assemblages using environmental DNA (eDNA) metabarcoding at 100 stations across three depth layers (300 samples) in Maizuru Bay, a semi-enclosed coastal bay in Japan. Concurrent hydrographic measurements were used to examine associations between fish assemblage composition and environmental conditions. We detected 216 fish species, with mean species richness decreasing from 28.4 at the surface to 16.4 at the bottom. Thirty-three species were detected exclusively below the surface. Fish assemblage composition differed significantly among depth layers (PERMANOVA *R²* = 0.119, *p* < 0.001) and exhibited pronounced horizontal turnover along the bay axis. Although depth, geographic position, and measured environmental variables were associated with assemblage composition, they collectively explained only 13.0% of the variation. The independent contribution of spatial position (2.9%) exceeded those of depth (0.8%) and measured environmental variables (0.9%). These findings demonstrate substantial three-dimensional heterogeneity in coastal fish assemblages that cannot be fully explained by conventional hydrographic measurements. Our study highlights the importance of integrating horizontal and vertical sampling to characterize coastal fish biodiversity and improve marine ecosystem monitoring.

**Highlights:** High-density eDNA surveys resolved three-dimensional coastal fish assemblages.

Fish species richness declined from surface to bottom waters.

Fish assemblages differed vertically and turned over horizontally across the bay.

Measured predictors explained only 13% of community variation.

Depth-resolved sampling improved the characterization of coastal fish diversity.

## 1. Introduction

Bays rank among the most valuable yet most degraded ecosystems on the planet (Lotze et al., 2006; Barbier et al., 2011). In coastal bays, freshwater inflow, tidal exchange, and local circulation create strong gradients in temperature, salinity, and turbidity, producing distinct water masses and vertical stratification (Officer, 1976; Yanagi, 1999). This oceanographic heterogeneity creates habitat mosaics over scales from hundreds of metres to kilometres, shaping the spatial organization of coastal communities (García-Charton et al., 2004; Roland Pitcher et al., 2012). Understanding how oceanographic heterogeneity structures fish communities is thus essential for predicting fish distributions, identifying ecologically important habitats, and designing spatial conservation and monitoring strategies in coastal ecosystems under changing environmental conditions (Worm et al., 2006; Embling et al., 2012; Moore et al., 2016). However, direct evidence linking fine-scale oceanographic features to whole-community organization remains limited, largely because obtaining spatially extensive community data at ecologically relevant spatial scales has been technically challenging (Ushio et al., 2023).

Among coastal organisms, fishes are particularly informative for testing this linkage: as highly mobile nekton, they can actively select among water masses and exploit the three-dimensional structure of the water column, and their distributions therefore respond directly to hydrographic gradients (Breitburg, 2002; Tang et al., 2026). Conventional approaches for investigating coastal fish communities, including trawl surveys, gill-net sampling, and underwater visual censuses, have demonstrated that fish communities often exhibit clear vertical and horizontal segregation associated with environmental gradients (e.g., Elliott & Hemingway, 2002; Able, 2005; Masuda, 2008; Reis-Filho et al., 2019; Freitas et al., 2025). Nevertheless, conventional surveys are generally labor-intensive, taxonomically selective, and often difficult to implement in structurally complex coastal environments. Obtaining spatially continuous information across multiple depth layers and extensive sampling stations is thus challenging; as a result, how oceanographic heterogeneity structures fish communities across both horizontal and vertical dimensions remains poorly understood.

Environmental DNA (eDNA) metabarcoding has rapidly emerged as a powerful tool that enables simultaneous, large-scale sampling of fish communities across spatial scales as large as an entire bay, encompassing multiple depth layers (Thomsen et al., 2016; Canals et al., 2021; Jeunen et al., 2020). That capacity is now being realised at national scale: programmes in Japan have assembled hundreds of coastal sites into datasets spanning the entire archipelago, each site represented by a single surface sample (Doi et al., 2026; Osada et al., 2026). By analyzing extra-organismal DNA fragments in water (i.e., eDNA), eDNA metabarcoding allows for inferring species composition without physically capturing animals (Deiner et al., 2017). Because eDNA metabarcoding requires only water sampling in the field, it can equally support high-density surveys across multiple depth layers and numerous sampling stations, providing a practical way to match the spatial scale of biological signals to oceanographic heterogeneity. Whether one such sample adequately represents its site depends on how much structure exists within it, which has not been quantified.

Maizuru Bay, a semi-closed bay on the Sea of Japan, offers a suitable system for this question. An earlier eDNA metabarcoding survey there detected 128 fish species from 47 stations in a single 6-h collection, recovering about 63% of the species recorded by 140 underwater visual censuses over 14 years for which reference sequences were available, as well as at least 23 species that the censuses had missed (Yamamoto et al., 2017). That survey also found community composition to be spatially autocorrelated only within about 800 m, suggesting that fish communities in this bay are structured at scales of several hundred metres. Here, we combined eDNA metabarcoding with concurrent oceanographic profiling to resolve the three-dimensional structure of the fish community in the bay. Sampling 100 stations across three depth layers (*n* = 300) and concurrently measuring temperature, salinity, turbidity, and chlorophyll a, we asked how fine-scale heterogeneity in oceanographic conditions shapes the three-dimensional spatial distribution of coastal fish communities. To address this, we specifically examined (i) how community composition differed among vertical depth layers, (ii) how it varied horizontally along the bay axis, and (iii) to what extent measured environmental variables explained these spatial patterns. By linking fine-scale biodiversity patterns to concurrently measured oceanographic structure, this study demonstrates the potential of spatially resolved eDNA surveys to reveal how oceanographic heterogeneity structures coastal fish communities. We further ask what the structure resolved here implies for the design and interpretation of the broad-scale assessments described above.

## Methods

### Field survey

Seawater sampling for fish eDNA metabarcoding was conducted in Maizuru Bay, Kyoto Prefecture, Japan (35°29′N, 135°23′E). This semi-closed bay has a surface area of approximately 22.87 km², a maximum depth of about 30 m, and is connected to the Sea of Japan through a narrow mouth on its northern side. This bay is largely separated into the west and east bays by Toshima Island and exchanges water only weakly with the open sea (Miwa & Ikeno 2007).

The water-sampling survey was conducted on 21 and 22 June 2016, at 100 stations arranged in a grid pattern with approximately 400 m spacing (**Fig. 1**). The average depth across the 100 stations was approximately 15 m. On both days, the survey proceeded from the bay mouth toward the inner bay. At each station, seawater was collected from three depth layers: surface, middle, and bottom. The middle layer was defined as 5 m below the surface, corresponding to just below the pycnocline, while the bottom layer was approximately 1 m above the seafloor. These eDNA samples used in this present study are the same as those described by Fukaya et al. (2021).

**Figure 1.**
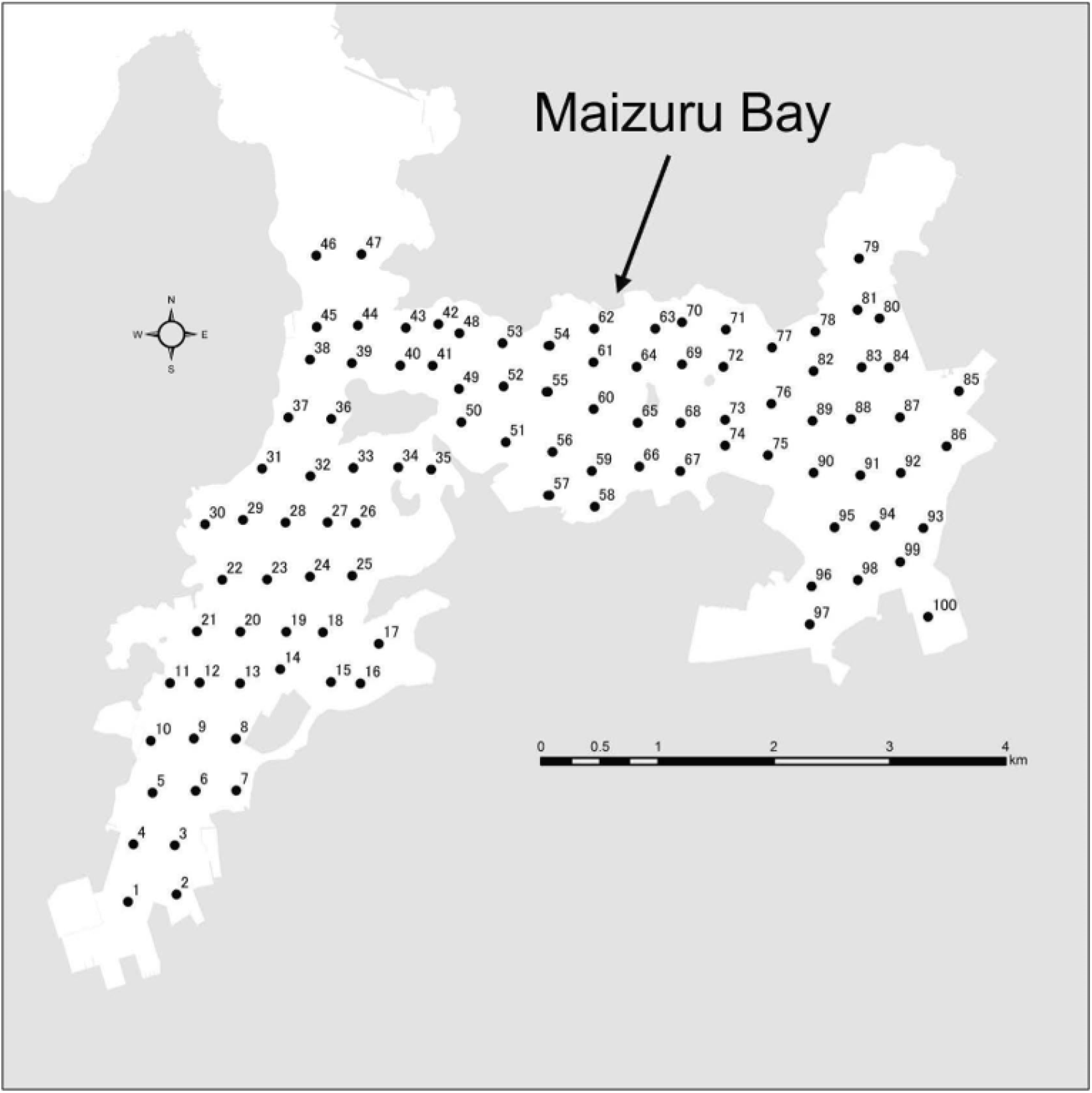
Survey stations for eDNA sampling in Maizuru Bay. Map showing the locations of 100 sampling stations in Maizuru Bay, Kyoto, Japan. Stations were arranged on an approximately 400 m grid covering the entire bay. Water samples were collected at three depth layers (surface, middle, and bottom) at each station.

Water samples were collected using a large ladle for surface water and vanDorn samplers for the middle and bottom layers. At each station and depth, one liter of water was collected into a 1.3-L plastic bottle, which had been pre-rinsed with a portion of the sample water. Immediately after collection, 1 mL of 10% benzalkonium chloride solution was added to each bottle and gently mixed to prevent DNA degradation (Yamanaka et al., 2017). The bottles were then stored in opaque containers to protect the samples from sunlight exposure. To minimize cross-contamination, disposable gloves were changed between sampling stations. During sampling, environmental variables—water temperature, salinity, turbidity (Formazin Turbidity Units), and chlorophyll a concentration—were measured using a CTD profiler (COMPACT-CTD: ASTD151, JFE Advantech, Hyogo, Japan) (**Table S1**).

### Water filtration and extraction

Water samples were filtered on the same day as the field survey using 47-mm glass fiber filters (GF/F; GE Healthcare Japan, Tokyo, Japan; nominal pore size: 0.7 μm) and an aspirator in the laboratory. Filters were folded with the filtration surface inward, wrapped in aluminum foil, and stored at -20°C until eDNA extraction. All procedures, from water sampling to filtration, were completed within seven hours. To prevent eDNA carryover, all filtration equipment was soaked in 0.1% sodium hypochlorite for at least five minutes, then thoroughly washed with tap water and rinsed with distilled water (DW) to remove residual bleach. As a filtration negative control to validate this decontamination process, one liter of DW was filtered after every 30 water samples using bleached equipment. A total of 10 filtration negative controls were prepared and subsequently processed through DNA extraction and PCR in the same manner as the environmental samples. DNA extraction from GF/F filters followed Yamamoto et al. (2016) using the DNeasy Blood and Tissue Kit (Qiagen, Hilden, Germany). Prior to adding the proteinase K (ProK) mix solution, the remaining water was centrifuged at 5,000 G for 3 min and collected in Salivette tubes (Sarstedt AG & Co. KG, Nümbrecht, Germany). The ProK mix solution (ProK 20 µL, Buffer AL 200 µL, Milli-Q water 200 µL) was then added directly to each filter. Finally, DNA was eluted with 100 µL of Buffer AE and extracted DNA solutions were stored at -20°C until further analysis.

### Paired-end library preparation for quantitative eDNA metabarcoding

Fish eDNA metabarcoding was performed using a quantitative MiSeq approach based on the MiFish metabarcoding method (Miya et al., 2015; Ushio et al., 2018). A two-step PCR protocol was used to amplify a fragment (163–185 bp) of the mitochondrial 12S rRNA gene and to add sample-specific index sequences. The first PCR was conducted in eight replicates per sample using the MiFish-U primer set. Each 12-μL reaction contained 6.0 μL of 2× KAPA HiFi HotStart ReadyMix (Kapa Biosystems, Wilmington, MA, USA), 1.4 μL of a 5 μM MiFish primer mixture, 1.0 μL of template DNA, 1.0 μL of an internal-standard DNA mixture, and 2.6 μL of Milli-Q water. The internal-standard mixture comprised three synthetic DNA sequences adjusted to 5, 25, and 50 copies/μL and was added to each reaction to enable sample-specific conversion of sequence reads to DNA copy numbers. The PCR conditions were set as follows: initial denaturation at 95°C for 3 min, followed by 40 cycles of denaturation at 98°C for 20 s, annealing at 65°C for 15 s, and extension at 72°C for 15 s, with a final extension at 72°C for 5 min. The first-PCR products were purified using AMPure XP magnetic beads (Beckman Coulter, Brea, CA, USA), and double-stranded DNA concentrations were measured using the QuantiFluor ONE dsDNA System (Promega, Madison, WI, USA). To assess the contamination on PCR steps, Milli-Q water 1µL is used for template as PCR negative control.

In the 2nd PCR, MiSeq adaptor and 8-bp index sequences were added to both ends of the amplicons (Hamady et al., 2008). To reduce variation in sequencing depth among samples, first-PCR products were diluted to 0.1 ng/μL before use as templates; products with concentrations below 0.1 ng/μL were used without dilution. Each 12-μL second-PCR mixture contained 6.0 μL of 2 × KAPA HiFi HotStart ReadyMix, 0.83 μL each of the forward and reverse Illumina XT index primers, 2.0 μL of the first-PCR product, and 2.34 μL of double-distilled water. Thermal cycling comprised initial denaturation at 95°C for 3 min, 12 cycles of 98°C for 20 s and 72°C for 15 s, and a final extension at 72°C for 5 min. Non-target amplification products were removed by size selection using the E-Gel SizeSelect system (Thermo Fisher Scientific, Waltham, MA, USA). The size distribution of the purified library was verified using a Bioanalyzer (Agilent Technologies, Santa Clara, CA, USA), with the target MiFish library producing an approximately 370-bp fragment including adapters. Library concentrations were measured using the QuantiFluor ONE dsDNA System, and the pooled library was adjusted to a final concentration of 11 pM. Paired-end sequencing was performed using an Illumina MiSeq platform with a MiSeq Reagent Kit v2.

### Bioinformatic analysis

Paired-end sequence reads were processed using the PMiFish pipeline version 2.4.1 with an ERS-custom workflow and USEARCH version 11. Forward and reverse reads were merged, primer sequences were removed, and reads were trimmed and quality-filtered. Sequences shorter than 50 bp were discarded. Error correction and denoising were performed, and putative chimeric and erroneous sequences were removed to generate amplicon sequence variants. Sequences representing less than 0.01% of the reads within a sample were excluded. The retained sequences were compared with the MiFish reference database dated 21 April 2022 using USEARCH global alignment. Taxonomic assignments were evaluated using sequence similarity to reference sequences, with searches conducted over an identity range of 85–100%. Representative sequences and their highest-ranking database matches were subsequently examined using BLAST. When multiple taxa showed equally high sequence similarity and could not be distinguished using the amplified region, sequences were assigned to the lowest taxonomic level supported by the reference data and information on the regional fish fauna.

The DNA concentration (copies/μL) was calculated from the sequence reads according to the method described by Ushio et al. (2018). Briefly, linear regression with an intercept set to zero was performed to examine the relationship between sequence reads and the copy numbers of the internal standard DNAs for each sample. Sequence reads of nonstandard fish DNA were converted to copy numbers by dividing the number of MiSeq sequence reads by the sample-specific regression slope. Each PCR-negative control corresponded to a portion of the sample, and copy numbers in the PCR-negative controls were subtracted from the corresponding samples to remove contaminants. Owing to the absence of research on the limit of quantification in quantitative eDNA metabarcoding, eDNA concentrations of less than one copy/μL were denoted as zero copies/μL.

### Statistical analysis

When multiple ASVs were assigned to the same species, their eDNA copy numbers were summed and used for subsequent statistical analyses. Although eDNA concentrations (copies/L) were estimated for each taxon by the quantitative MiSeq method, all community analyses were based on presence/absence data. Quantitative metabarcoding data remain only semi-quantitative for relative species abundance because amplification efficiency differs among taxa and primer-template combinations. Presence/absence data are therefore more robust for comparing community composition among samples. Taxa not detected in any sample were excluded, leaving 216 fish species across 300 samples (100 stations × 3 depth layers).

Because bathymetry varies across the bay, the bottom layer did not correspond to a fixed isobath: actual sampling depths ranged from 6.8 to 27.5 m (median = 14.0 m). Sampling depth in this layer was uncorrelated with longitude (Spearman ρ = -0.17, *p* = 0.083) but positively correlated with latitude (ρ = 0.58, *p* < 0.001). This covariance is taken into account when interpreting the latitudinal term below.

Species richness was calculated as the number of species detected per sample and projected onto a map of Maizuru Bay, overlaid with coastline shapefiles (JGD2000, EPSG:4612, transformed to WGS84) and drawn separately for each depth layer. Because the three layers were sampled at the same stations, richness values were treated as repeated measures: differences among layers were tested with a Friedman rank sum test blocked by station, followed by pairwise Wilcoxon signed-rank tests with Bonferroni correction. Horizontal gradients were assessed by Spearman rank correlation between richness and longitude within each layer. Taxonomic overlap among layers was visualised with a Venn diagram.

To investigate community dissimilarity among samples, we calculated the Sørensen dissimilarity between all pairs of samples. NMDS was performed with the metaMDS function (*k* = 3, trymax = 500, maxit = 999); stress values and convergence were recorded. To visualize the spatial structure of community composition, NMDS scores were converted to a continuous color scale in HCL (hue-chroma-luminance) space: NMDS1 was rescaled to hue (200 to 360) and NMDS2 to lightness (40 to 90), with chroma fixed at 60, yielding a unique color for each site. Because the sign of NMDS axes is arbitrary, axes were reflected so that NMDS1 correlated positively with longitude and NMDS2 with latitude; reflection is a mirror operation and does not alter the ordination. Colored points were plotted on a map of the bay separately for each depth layer, allowing intuitive interpretation of spatial turnover in fish communities. NMDS was additionally run within each layer separately; because these layer-wise ordinations were rescaled independently, colors are not comparable across panels.

Differences in community composition among depth layers were tested by permutational multivariate analysis of variance (PERMANOVA; adonis2). Because the three layers were sampled at the same stations, samples were not independent, and permutations were therefore restricted to within stations (9999 permutations). Pairwise comparisons between layers used the same restricted design with Bonferroni correction. Homogeneity of multivariate dispersions among layers was tested with betadisper and permutest (9999 permutations) to determine whether any PERMANOVA result reflected a shift in group centroids rather than a difference in within-group dispersion. Because the bottom layer spanned a range of sampling depths, we further tested, within that layer alone, whether actual sampling depth explained community composition, using PERMANOVA with depth, longitude, and latitude as marginal terms (9999 permutations).

To evaluate the influence of environmental variables on community composition, we performed distance-based redundancy analysis (dbRDA) using the dbrda function, applying a square-root transformation of the dissimilarities to avoid negative eigenvalues arising from the non-Euclidean distance measure. Candidate predictors included depth layer (categorical), environmental variables measured by the CTD profiler (temperature, salinity, turbidity, and chlorophyll a), and geographic coordinates (longitude and latitude) to account for spatial gradients. Turbidity and chlorophyll a were strongly right-skewed and were natural-log-transformed before standardisation; temperature and salinity were used untransformed. All continuous predictors were standardised to zero mean and unit variance prior to analysis. Predictors were selected by forward selection based on adjusted *R²* (ordiR2step, 999 permutations). Model significance, the marginal effect of each retained predictor, and the significance of individual constrained axes were assessed using permutation tests with 9999 replicates, and variance inflation factors (VIFs) were inspected to check for collinearity. The ordination biplot shows sites, vectors for the continuous predictors, and centroids for the categorical depth-layer term, facilitating interpretation of the environmental gradients shaping the fish community. Because depth and temperature proved strongly collinear as a consequence of vertical stratification, the explained variation was additionally partitioned among three predictor sets (depth layer, environmental variables, and space defined by longitude and latitude) using varpart. Note that all candidate predictors were included in this partitioning, irrespective of whether they were retained by forward selection in the dbRDA. All analyses were performed in R v4.6.1 (R Core Team, 2026) using the vegan package v2.7-5 (Oksanen et al., 2026), with a fixed random seed for all stochastic procedures.

## Results

### The oceanographic structure in Maizuru Bay

We first clarified vertical stratification and horizontal differences in water-mass structure in Maizuru Bay using in situ oceanographic data from a conductivity, temperature, and depth profiler (CTD). A thermocline was consistently observed at approximately 5 m depth, based on the vertical profiles of water temperature (**Fig. S1**). Surface waters were generally warmer than the middle and bottom layers across the bay, indicating that thermal stratification was well-developed during the survey period. In addition to the vertical structure, oceanographic conditions also varied horizontally between the western and eastern parts of the bay. At the surface, higher temperatures and lower salinity were observed in the west bay and in river-influenced areas of the inner east bay (**Fig. S2**). The same west–east difference in temperature was evident in the middle and bottom layers, where temperatures were lower in the east bay, with particularly cold bottom water occurring in the northwestern part of the east bay. Salinity, however, varied only narrowly across the bay (29.9–34.2), remaining close to full-strength seawater throughout. These oceanographic features indicate differences in water mass structure between the west and east bays. Based on these oceanographic patterns, particularly the spatial distribution of water temperature, we delineated the bay into western and eastern sectors using a north–south transect passing through stations 48–50, with stations 1–50 assigned to the west bay and stations 51–100 to the east bay. This classification was used in subsequent analyses of eDNA-based fish community structure.

### eDNA metabarcoding revealed fish communities driven by the oceanographic structure

A total of 216 fish species were detected across the 300 samples. In the filtration negative control, 1.05 copies of *Trachurus japonicus* were detected, while no fish DNA was detected in the PCR negative control (**Table S4**). Species richness per sample ranged from 1 to 64 and decreased monotonically with depth: 28.4 ± 12.0 species (mean ± standard deviation) in the surface layer, 20.5 ± 8.9 in the middle layer, and 16.4 ± 7.8 in the bottom layer (**Fig. S3**). The difference among layers was significant (Friedman χ² = 83.4, df = 2, *p* < 0.001), and all pairwise comparisons remained significant after Bonferroni correction (surface vs middle, *p* < 0.001; surface vs bottom, *p* < 0.001; middle vs bottom, *p* < 0.001). Richness also varied horizontally, decreasing from west to east in all three layers, although the gradient weakened with depth (Spearman ρ = - 0.32, *p* = 0.001 in the surface layer; ρ = -0.33, *p* = 0.001 in the middle layer; ρ = -0.22, *p* = 0.030 in the bottom layer). Within the bottom layer, richness declined only weakly with sampling depth, a marginal trend that explained little variation (ρ = -0.19, p = 0.055). The lower richness of the bottom layer therefore cannot be attributed simply to sampling deeper water at some stations.

The number of species detected per layer followed the same order as sample-level richness: 171 species in the surface layer, 144 in the middle layer, and 123 in the bottom layer. Of the 216 species, 86 (40%) were detected in all three layers (**Fig. 2**). Species restricted to a single layer numbered 47 (22%) in the surface layer, 19 (9%) in the middle layer, and 14 (6%) in the bottom layer. Species shared by exactly two layers were less frequent: 27 (12%) between the surface and middle layers, 12 (6%) between the middle and bottom layers, and 11 (5%) between the surface and bottom layers. The smallest two-layer overlap was between the surface and bottom layers, that is, between the two most vertically distant layers.

**Figure 2.**
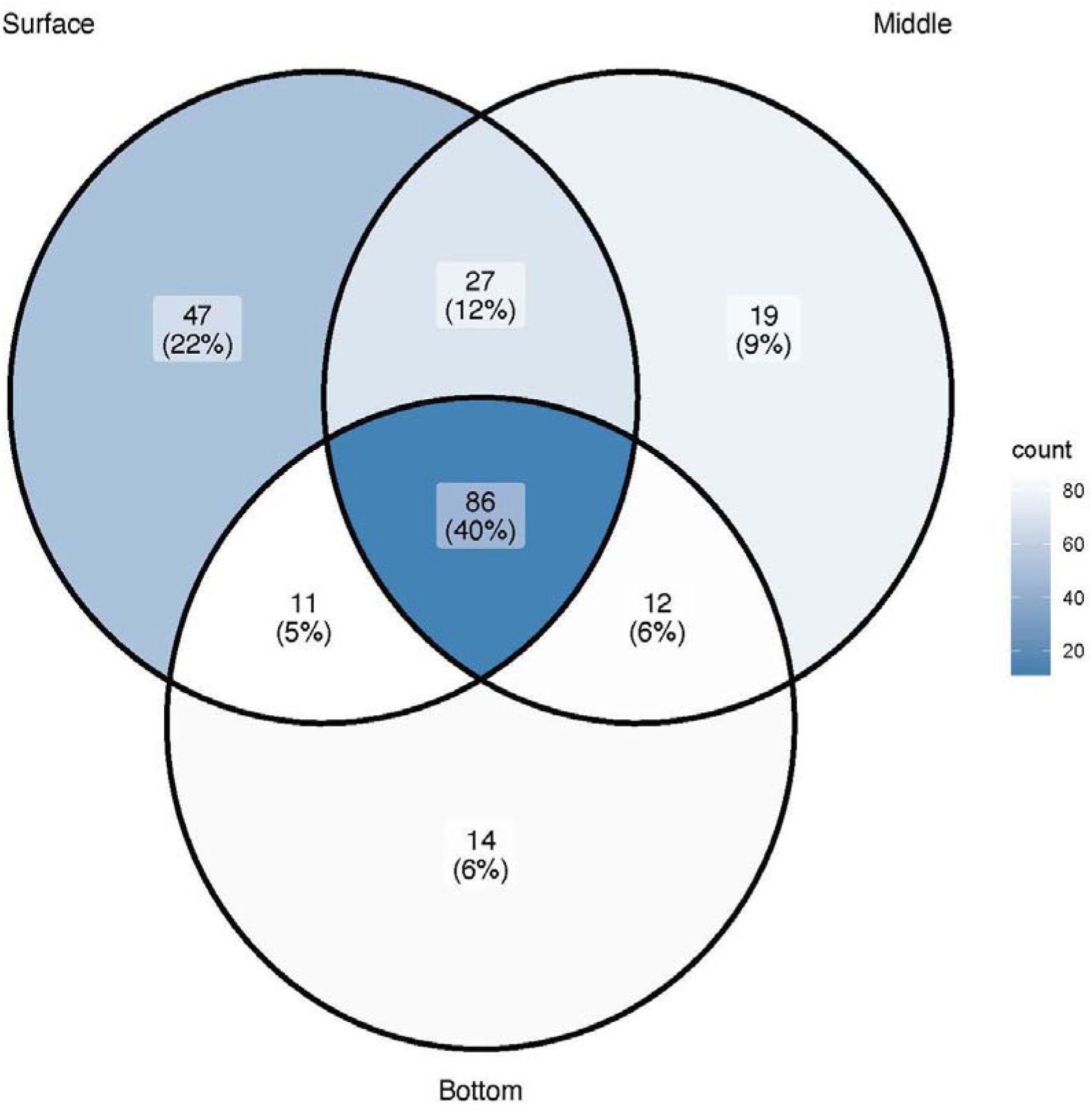
Species overlap among depth layers. Venn diagram showing the number of fish species detected across three depth layers (surface, middle, and bottom). Numbers indicate species shared among layers or unique to each layer, with percentages shown in parentheses.

The species detected in each layer were ecologically consistent with the vertical pattern. The most frequently detected species were common in the bay and were found throughout the water column: *Engraulis japonicus* (98% of samples), *Trachurus japonicus* (92%), *Acanthopagrus schlegelii* (91%), *Nuchequula nuchalis* (89%), and *Konosirus punctatus* (84%). Species with a strong vertical bias, by contrast, corresponded to their known habitats. Surface-associated species were detected mainly at the surface (*Mugil cephalus*, 97 versus 34 stations at the surface and bottom; *Hyporhamphus sajori*, 55 versus 4), whereas demersal gobies and dragonets were detected mainly at the bottom (*Acentrogobius virgatulus*, 26 versus 81; *Amblychaeturichthys hexanema*, 0 versus 46; *Callionymus valenciennei*, 8 versus 41). Species detected only at the bottom included several benthic taxa, such as the gobies *Amblychaeturichthys sciistius* and *Paratrypauchen microcephalus*, whereas those detected only at the surface included freshwater and estuarine species such as *Gnathopogon elongatus*, *Odontobutis obscura*, and *Eutaeniichthys gilli*.

Non-metric multidimensional scaling (NMDS) on Sørensen dissimilarities converged with a stress of 0.168 (*k* = 3); layer-wise ordinations also converged (stress = 0.153, 0.166, and 0.161 for the surface, middle, and bottom layers, respectively). Mapping NMDS scores onto the bay revealed a west-to-east turnover in community composition across all three layers (**Fig. 3**).

**Figure 3.**
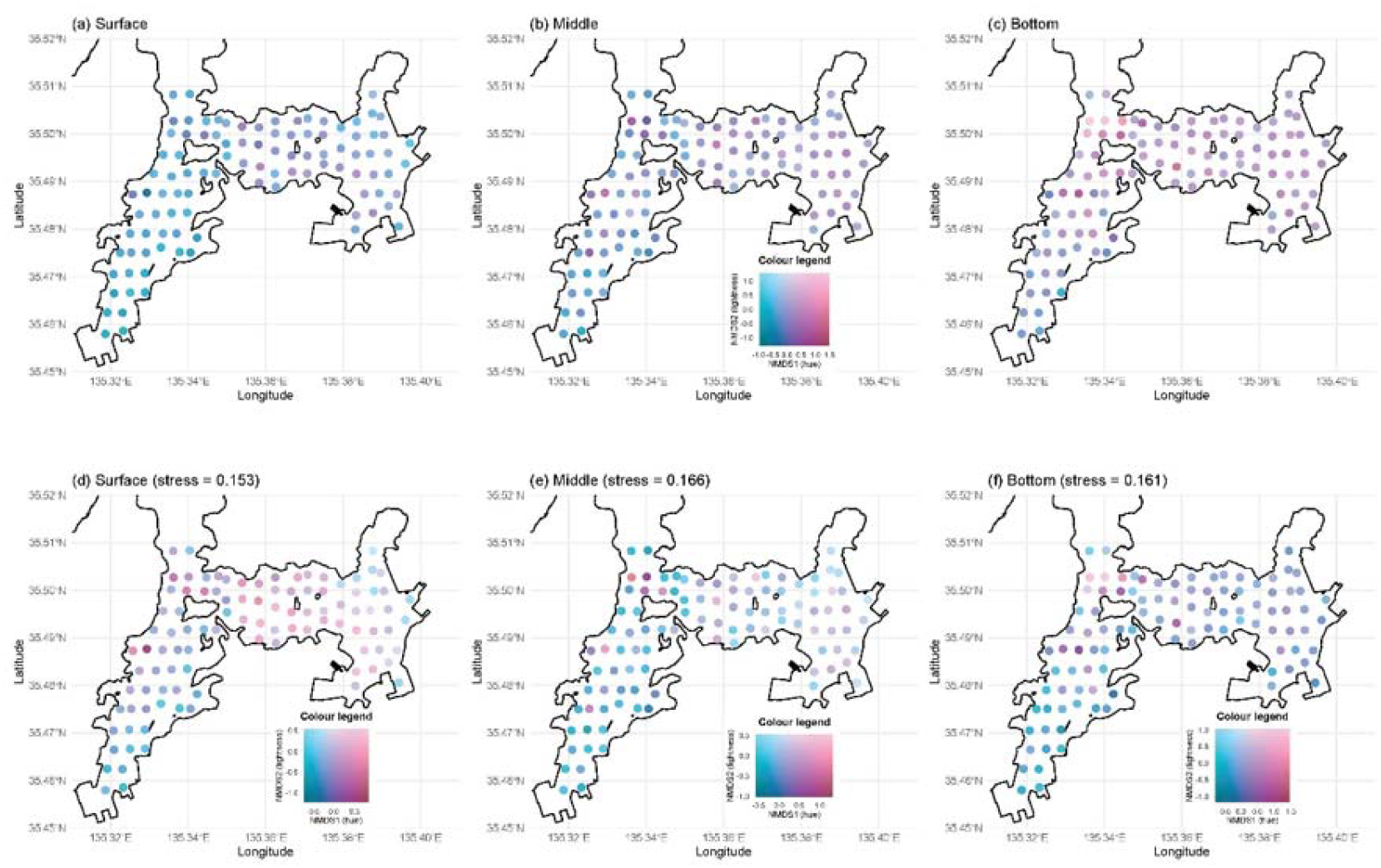
Spatial patterns of fish community composition based on NMDS. NMDS scores mapped onto Maizuru Bay, with NMDS1 represented as hue and NMDS2 as lightness in HCL colour space. Panels (a–c) show scores from a single ordination of all samples, mapped separately for the surface, middle, and bottom layers; panels (d–f) show ordinations run independently within each layer. Colours are comparable among panels (a–c), which derive from a single ordination, but not among (d–f) or between the two sets, which were rescaled independently.

Community composition differed significantly among depth layers (PERMANOVA with permutations restricted within stations: *R²* = 0.119, *F* = 19.99, *p* < 0.001), and all pairwise contrasts were significant after Bonferroni correction (*p* < 0.001). The difference was largest between the surface and bottom layers (*R²* = 0.157) and smallest between the surface and middle layers (*R²* = 0.045), with the middle-bottom contrast intermediate (*R²* = 0.064). Multivariate dispersion did not differ among layers (betadisper: *F* = 0.49, *p* = 0.609; mean distance to centroid = 0.366, 0.377, and 0.379 for the surface, middle, and bottom layers). The PERMANOVA result can therefore be attributed to a shift in the centroid of community composition rather than to differences in within-layer heterogeneity. Within the bottom layer, actual sampling depth had a small but significant effect on composition (*R²* = 0.020, *p* = 0.012), comparable in magnitude to that of longitude (*R²* = 0.028, *p* = 0.001), whereas latitude was not significant once sampling depth was accounted for (*R²* = 0.008, *p* = 0.490).

Forward selection in distance-based redundancy analysis (dbRDA) retained depth layer, longitude, latitude, chlorophyll a, salinity, and temperature (adjusted *R²* = 0.130); only turbidity was not retained (**Table 1**). The final model was significant (*F* = 7.40, *p* < 0.001), as were the first five constrained axes (*p* < 0.05), and the first two axes explained 7.9% and 3.5% of the total variation, respectively (**Fig. 4)**. Ranked by marginal F, longitude was the strongest single predictor, followed by latitude, chlorophyll a, and depth layer, whereas salinity and temperature had comparatively weak effects (**Table 1**). Samples were arranged along dbRDA1 according to longitude, while the centroids of the three depth layers separated along dbRDA2. Turbidity, the only variable not retained by forward selection, was nonetheless significantly correlated with the constrained ordination when fitted post hoc (envfit *R²* = 0.36, *p* < 0.001; **Fig. S4**).

**Figure 4.**
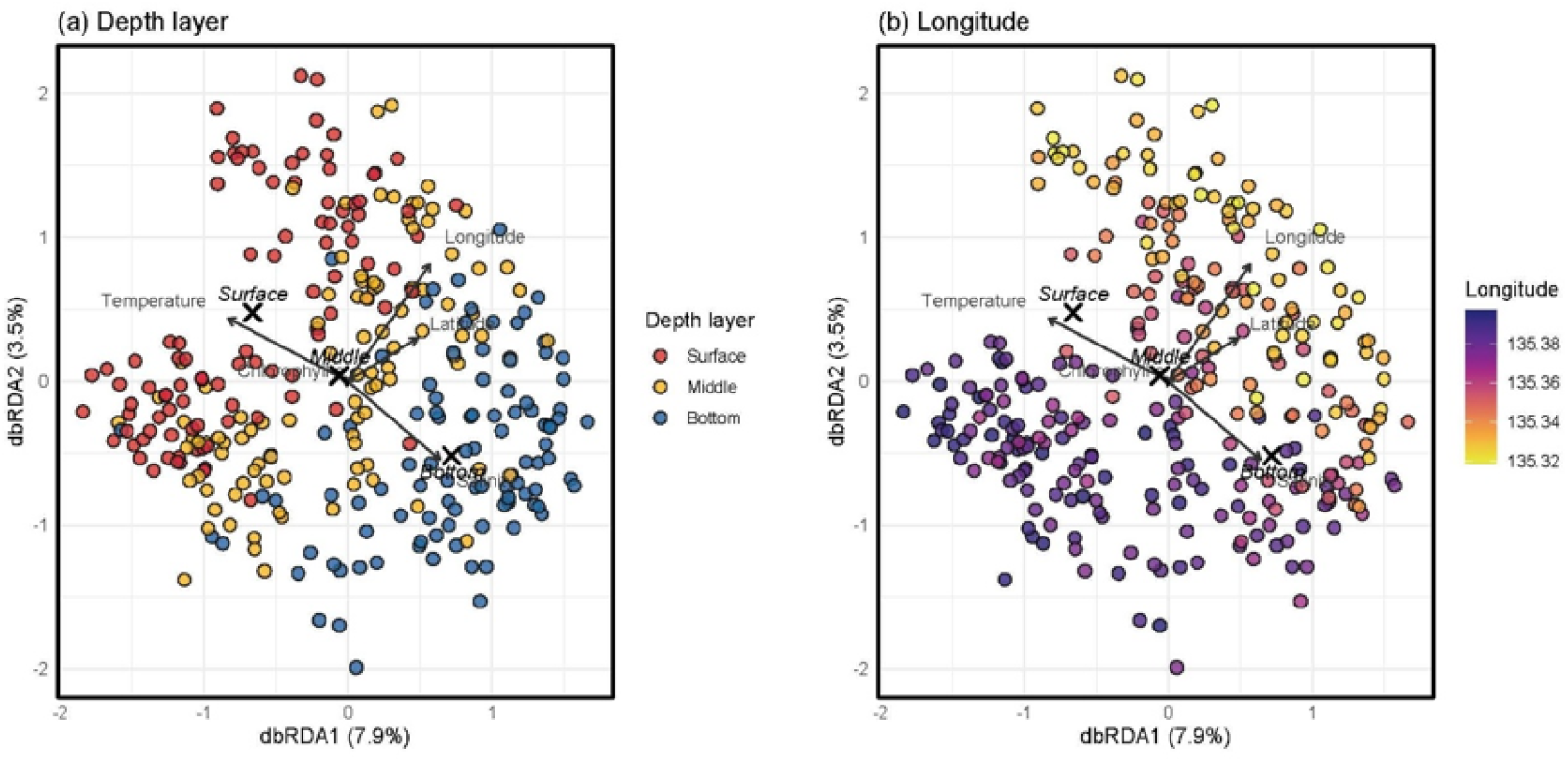
dbRDA of fish community composition in Maizuru Bay, based on Sørensen dissimilarities. The same ordination is shown twice, with samples coloured by (a) depth layer and (b) longitude. Arrows indicate the continuous predictors retained in the final model, and crosses mark the centroids of the three depth layers. Axis labels give the proportion of the total variation explained by each axis. The final model (depth layer + longitude + latitude + chlorophyll a + salinity + temperature) was significant (F = 7.40, *p* < 0.001; adjusted *R²* = 0.130; see **Table 1**). Axis orientation was fixed for reproducibility and does not affect interpretation.

**Table 1.** Results of distance-based redundancy analysis (dbRDA) of fish community composition in Maizuru Bay.

| **Predictor** | ***df*** | ***F*** | ***p*** | **Cumulative adj. *R<sup>2</sup>*** |
|---|---|---|---|---|
| Depth layer | 2 | 2.68 | < 0.001 | 0.06 |
| Longitude | 1 | 8.63 | < 0.001 | 0.107 |
| Latitude | 1 | 3.78 | < 0.001 | 0.122 |
| Chlorophyll a | 1 | 2.97 | < 0.001 | 0.126 |
| Salinity | 1 | 1.73 | 0.009 | 0.129 |
| Temperature | 1 | 1.40 | 0.055 | 0.130 |
| Residual | 292 |  |  |  |

Predictors are listed in the order in which they entered the model by forward selection based on adjusted *R<sup>2</sup>*; turbidity was not retained. F and p are from marginal permutation tests (9999 permutations) on the final model. Cumulative adj. *R<sup>2</sup>* is the value reached at each step of forward selection. Turbidity and chlorophyll a were log-transformed before analysis. Global model: F = 7.40, *p* < 0.001, adjusted *R<sup>2</sup>* = 0.130.

Temperature and the depth-layer term were strongly collinear (VIFs of 30.3 for temperature and up to 21.3 for depth layer), reflecting vertical thermal stratification. Rather than removing either term, both of which are of direct ecological interest, we used variation partitioning to separate their contributions. All predictor sets together explained 13.0% of the variation in community composition. Depth alone accounted for 0.8%, the environmental variables alone for 0.9%, and space (longitude and latitude) alone for 2.9%, while the largest single fraction (5.3%) was shared between depth and the environmental variables (**Fig. S5**). Turbidity showed the same pattern: correlated with the ordination but adding no explanatory power beyond the retained predictors. Although depth was thus strongly associated with community composition, its effect was largely inseparable from the vertical gradients in temperature and salinity that define stratification, and the largest independent contribution came from horizontal position within the bay.

## Discussion

Our high-density eDNA metabarcoding surveys (100 stations × 3 depth layers) revealed fish communities that were clearly organized along both vertical and horizontal axes. Although vertical differentiation was robust, horizontal position within the bay was at least as strong a correlate of community composition as depth. Several oceanographic variables were associated with community composition; together with depth and spatial position, however, they explained only a small fraction of the total variation. This indicates that the spatial organization of fish communities in this bay cannot be reduced to the environmental variables we measured.

### Vertical structure and the surface excess

Species richness declined monotonically from surface (28.4 ± 12.0) through middle (20.5 ± 8.9) to bottom (16.4 ± 7.8). The asymmetry extended to species overlap: 47 species were detected only in surface water, but only 14 at the bottom, while 86 occurred in all three layers. The bottom assemblage was thus held largely to be a depauperate subset of the shallower fauna rather than a distinct fauna of bottom specialists. Many coastal fishes have a pelagic larval stage before recruiting to their species-specific juvenile and adult habitats (Leis & Carson-Ewart, 2000). Because our survey was conducted in June, when larval fish abundance is generally highest in Japanese coastal waters (Kanou et al., 2002), pelagic larvae of numerous species were likely present in the surface waters. Recent studies have demonstrated that fish larvae release sufficient amounts of eDNA for molecular detection (Maruyama et al., 2014; Takeuchi et al., 2019). Consequently, surface-water eDNA likely included signals from both pelagic larvae and resident adult fishes, resulting in higher species richness than in deeper layers. Consistent with this, Yamamoto et al. (2017) detected demersal gobiid species in surface water of this bay and attributed these detections to their pelagic larvae, which are present in the bay during this season. The thermocline at around 5 metres may have further contributed to differences in eDNA signals between layers (Jeunen et al., 2020; Littlefair et al., 2021). Unlike conventional sampling methods, which target specific life stages, eDNA metabarcoding can detect genetic material from multiple developmental stages, potentially providing a broader perspective on fish community structure.

### Horizontal structure and the weak environmental signal

Longitude was the strongest predictor in the dbRDA, and variation partitioning confirmed that the pure spatial component exceeded the pure depth and environmental components. The small pure depth fraction reflects the sharing of most of the depth signal with temperature rather than a weak depth effect. Part of the longitudinal signal, too, may reflect a horizontal temperature gradient, since temperature was strongly correlated with longitude within the middle and bottom layers (Spearman ρ = -0.85 and -0.55; **Fig. S2**). Yet spatial position explained variation beyond that accounted for by the measured environmental variables, indicating that longitude was not merely a proxy for temperature. It may also reflect sub-basin identity: Toshima Island divides the bay into two weakly connected basins that differ in water exchange with the open sea and in terrestrial inputs (Miwa & Ikeno, 2007), neither of which our CTD variables captured. Latitude, increasing towards the bay mouth, may capture the influx of marine species from the Sea of Japan.

Among the four oceanographic variables, chlorophyll a had the strongest effect on community composition, though it ranked below the spatial predictors overall; salinity had a comparatively weak effect, and turbidity was not retained. That salinity mattered so little, despite its central role elsewhere, is itself informative. In estuaries, salinity is regarded as the primary determinant of fish distribution, and much of the conceptual framework for coastal fish ecology rests on this premise (Elliott & Whitfield, 2011; Whitfield, 2021). Maizuru Bay, however, receives limited freshwater input, and salinity varied only narrowly (29.9–34.2; **Fig. S2**). The steep gradient that structures estuarine assemblages is thus essentially absent, and the weak salinity effect we detected likely reflects this narrow range. Frameworks developed for estuaries may therefore transfer poorly to semi-closed bays with limited freshwater input, such as Maizuru Bay. More broadly, the environmental variables we measured explained only a small part of the community variation, suggesting that unmeasured factors may matter as much as those we recorded.

### Unmeasured drivers

Measuring additional variables not obtained in this study would be key to better understanding differences in fish community structure in the horizontal and vertical directions within the studied bay. First, although we did not measure dissolved oxygen (DO) as our CTD carried no oxygen sensor, DO is likely to be a key factor in explaining the vertical differentiation of fish communities. DO typically exhibits a steep vertical gradient; while it is high in the surface layer due to atmospheric exchange and photosynthesis, the bottom layer is prone to hypoxia through organic matter decomposition and stratification (Epifanio et al., 1983; Miwa & Ikeno, 2007). In semi-closed bays such as the one studied here, DO levels often drop significantly in the bottom layer during certain seasons. In addition to water temperature and chlorophyll a, such DO gradients may also have contributed to the vertical differentiation of fish communities. For example, while fish generally actively avoid hypoxic environments, tolerance to hypoxia varies between species (Breitburg, 2002; Zhang et al., 2019).

Second, benthic habitat, including substrate type, submerged vegetation, and structural complexity, may support a high degree of horizontal heterogeneity in fish communities, particularly in the bottom layer. Demersal fish, in particular, are highly dependent on the bottom substrate, hiding places, and feeding grounds (Caddy, 2008; Elliott et al., 2007), which may have had a significant influence on the formation and spatial differentiation of their assemblages. Because we did not characterise benthic habitat, its omission may have partially accounted for the large residual variation, particularly in the bottom layer.

Third, fish distributions may also be shaped by food resources, such as zooplankton and other prey. In coastal ecosystems, these food resources often concentrate near current boundaries, thermoclines, and river estuaries, where nutrients and organic matter accumulate (Hickey et al., 2010). Chlorophyll a, retained as a predictor in our model, is consistent with such a productivity-driven effect. The weak, marginal effect of temperature, too, may have been partly indirect and bottom-up, mediated through the distribution of food resources rather than through direct physiological responses (e.g., Möllmann et al., 2005).

Fourth, and most importantly, the strong spatial signal should be explained mechanistically. Maizuru Bay is characterised by partial isolation between its western and eastern areas, likely resulting in weak water exchange and long residence times. Quantifying distance from the bay mouth, water residence time, or other measures of hydrodynamic connectivity, ideally with a circulation model of the kind developed for this bay (Miwa & Ikeno, 2007), would help translate geographic coordinates into ecological processes. This would clarify whether the weak, compartmentalised circulation of the bay is what maintains the community differentiation we observed among its different parts. Such connectivity governs the transport not only of fish and their eggs, larvae, and prey, but also of eDNA itself, so that the location of an eDNA signal need not coincide with where an organism lives (Andruszkiewicz et al., 2019; Kasai et al., 2021). We therefore interpret longitude as a partial proxy for hydrodynamic connectivity, an interpretation that should be tested more explicitly in future eDNA-based studies of fine-scale fish community structure.

### Conservation implications

These results have practical consequences for broad-scale eDNA assessment of coastal biodiversity. In Japan, national eDNA programmes have invested in spatial extent, with 528 sites along more than 19,000 km of coastline at a spacing of tens of kilometres (Osada et al., 2026), and in temporal frequency, with 211 sites sampled as often as biweekly over more than a year (Doi et al., 2026), but both collect only surface water at a single location per site. Our results indicate what that leaves unmeasured. Within a bay of only 22.87 km², composition turned over horizontally in all three layers, and 33 of the 216 species were detected only below the surface; these were disproportionately demersal, the group most exposed to bottom hypoxia, loss of benthic habitat and trawling. The two scales agree on the cause: nationally, measured temperature and salinity explained only 1.7% of the variance in species co-occurrence (Osada et al., 2026), and here too the measured variables explained little while horizontal position carried more independent information than the environment. Where one surface sample stands for an entire semi-closed system, the resulting bias is structured rather than random: demersal species are systematically under-recorded and internal turnover is invisible. Because broad-scale eDNA assessments increasingly inform the delimitation of biogeographic boundaries and the setting of conservation priorities, that bias propagates into the management decisions they support, and needs to be allowed for in their design and interpretation.

### Conclusions

In conclusion, high-density eDNA metabarcoding resolved the three-dimensional structure of a coastal fish community across an entire bay. Community composition differed among depth layers, with vertical differentiation closely tied to thermal stratification, and turned over horizontally along the bay axis. The measured environmental variables explained only a small part of this variation, and horizontal structure was governed more by geographic position than by any single water-column variable, suggesting hydrodynamic processes our sampling did not capture. By matching the spatial resolution of biological sampling to that of oceanographic heterogeneity, spatially resolved eDNA surveys offer a practical route to linking biodiversity patterns with the physical processes that shape coastal systems.

## Acknowledgments

We sincerely thank Professor Yoh Yamashita, Associate Professor Keita W. Suzuki, Dr. Hideki Sawada, Ms. Misaki Shiomi, and the staff and students of Maizuru Fisheries Research Station, Kyoto University, Professor Michio Kondoh (Tohoku University), Professor Toshifumi Minamoto (Kobe University), and all members of this survey for logistical support during the field surveys in Maizuru Bay.

## Declaration of generative AI use

During the preparation of this manuscript, the authors used ChatGPT (OpenAI) to improve the English language and readability of the manuscript. The authors reviewed and edited all AI-assisted content and take full responsibility for the final content of the manuscript.

## CRediT authorship contribution statement

H. M. designed and performed research and wrote the manuscript; R. N. analyzed data and wrote the manuscript; T. S. J., Q. W. D., K. T. and R. M. contributed to conceptual discussions. All authors discussed the results, commented on the manuscript, and approved the final version.

## Data Accessibility Statement

The species-by-sample detection matrix for the 216 fish species and 300 samples, the accompanying CTD measurements of water temperature, salinity, turbidity and chlorophyll a, and the R code used for all community analyses and to generate the figures are available at figshare (https://figshare.com/s/47406c8e060a8b3657b8).

