## Supplementary material for "High-density environmental DNA metabarcoding reveals fine-scale three-dimensional structure of a fish community in a semi-closed bay": Figure S1-S5

**Supplementary information**

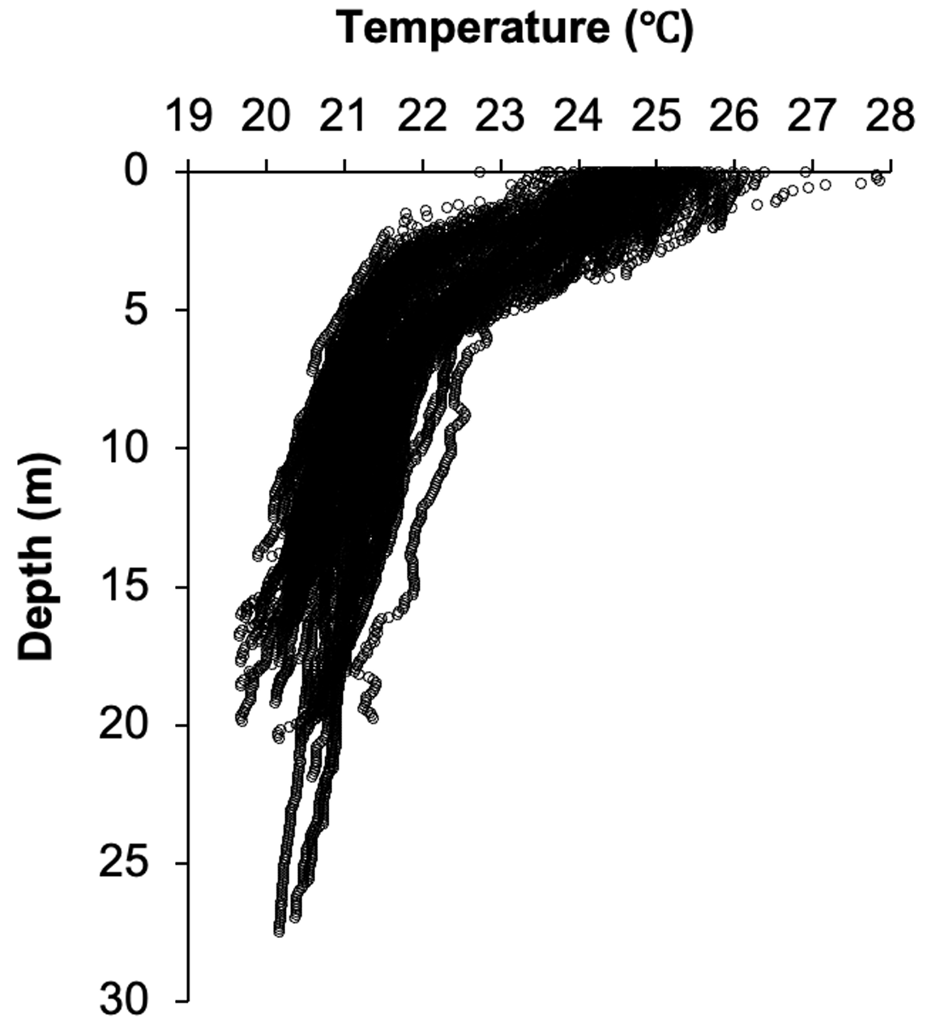

**Figure S1. Vertical profiles of water temperature in the west and east bays of Maizuru Bay.** Vertical profiles of water temperature in the west and east bays of Maizuru Bay. Profiles measured by CTD at the 100 sampling stations are shown. Each series is a single cast. A thermocline is present at approximately 5 m across the bay.

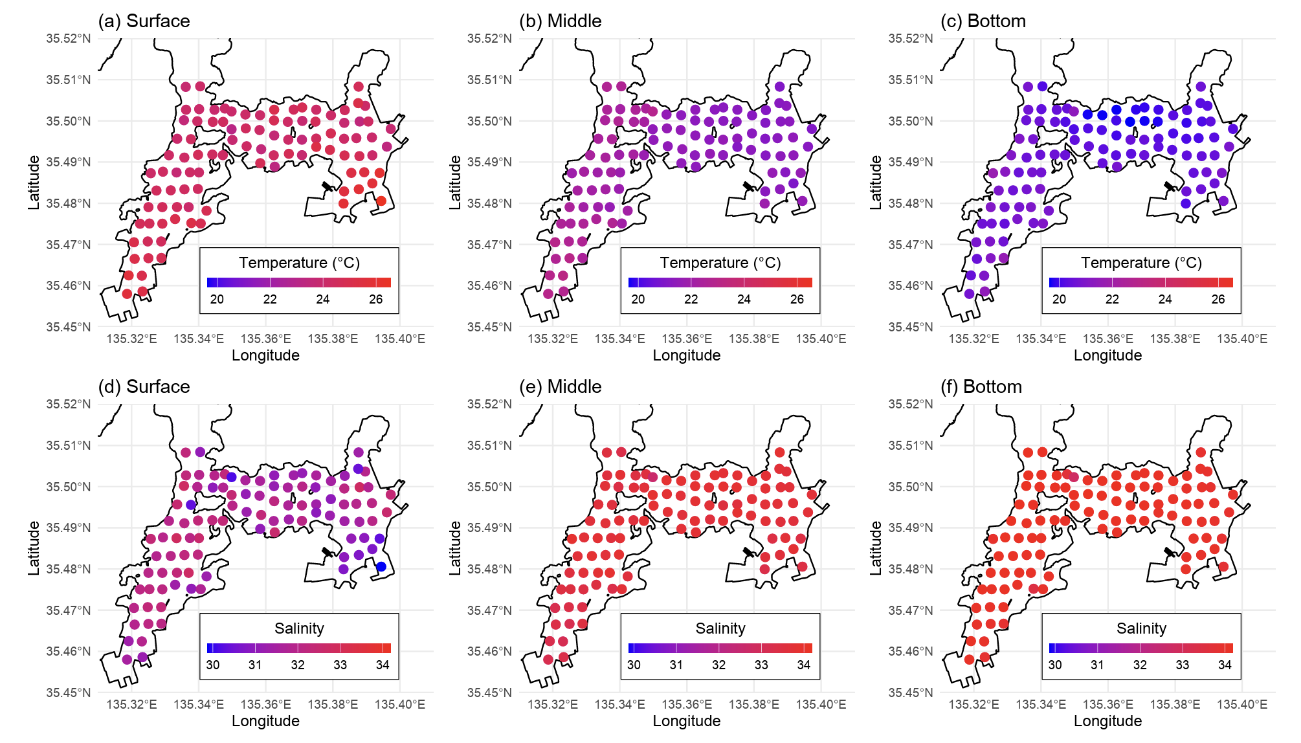

**Figure S2.** **Spatial distribution of temperature and salinity.** Water temperature (a–c) and salinity (d–f) at each sampling station for the surface, middle, and bottom layers. Colour scales are shared across layers within each variable.

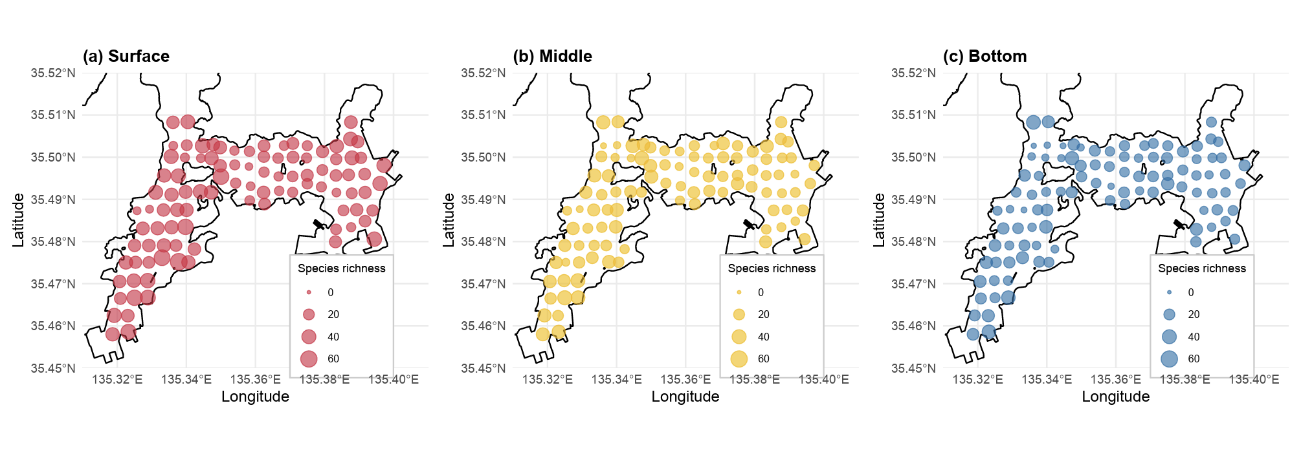

**Figure S3. Species richness at each sampling station for the (a) surface, (b) middle and (c) bottom layers.** Symbol size is proportional to the number of fish species detected per sample, on a scale shared across the three panels; colour distinguishes the layers and carries no additional information.

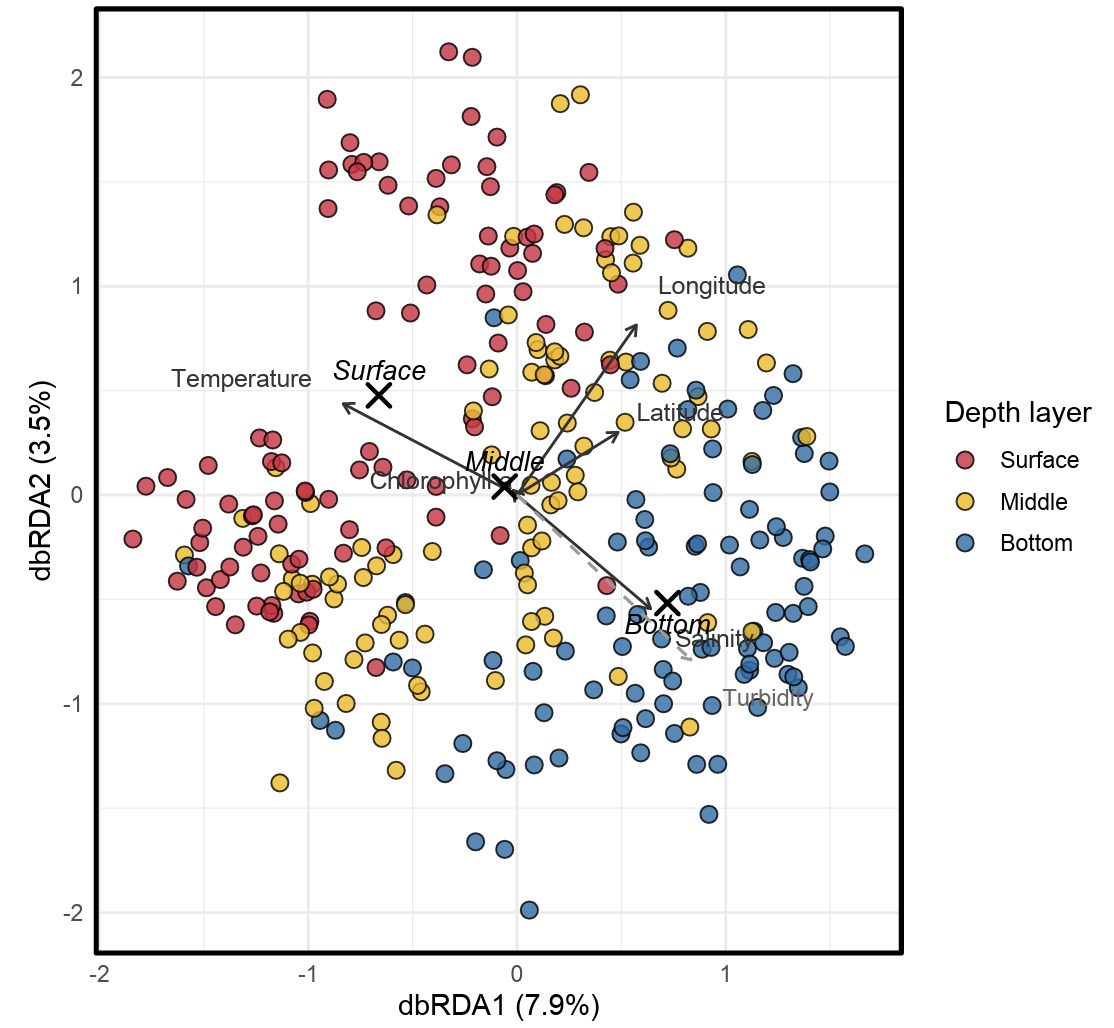

**Figure S4. Post hoc fit of turbidity onto the dbRDA ordination.** dbRDA of fish community composition, showing the predictors retained in the final model by forward selection (black solid arrows) together with turbidity, the only variable not retained. Turbidity was fitted post hoc onto the final ordination using envfit (grey dashed arrow; R² = 0.36, *p* < 0.001) and is rescaled for display. Samples are coloured by depth layer, and crosses mark the centroids of the three depth layers.

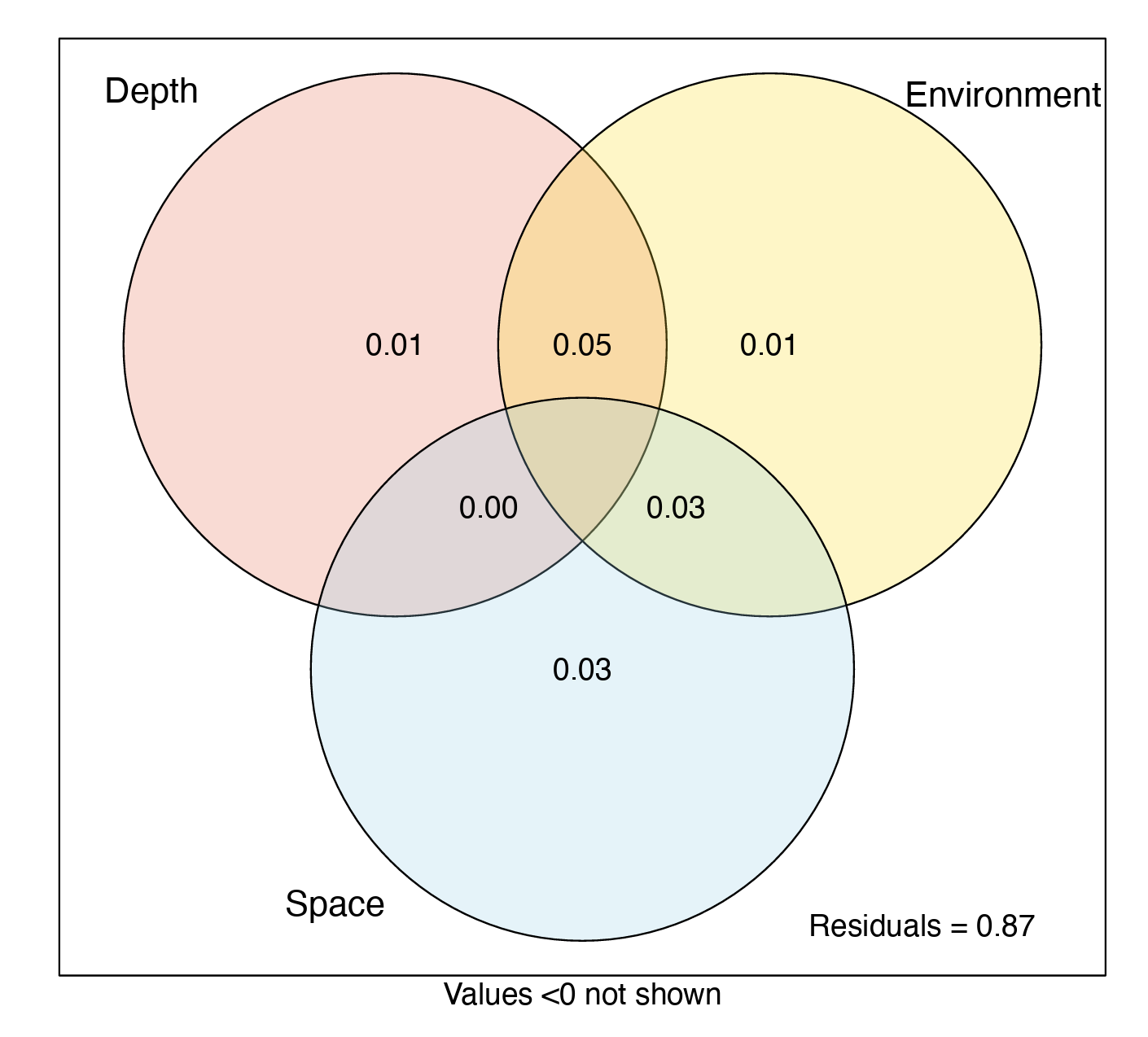

**Figure S5. Variation partitioning among depth, environment and space.** Variation partitioning of fish community composition among three sets of predictors: depth layer, environmental variables (temperature, salinity, turbidity and chlorophyll a), and space (longitude and latitude). All candidate predictors were included, irrespective of whether they were retained by forward selection in the dbRDA. Values are adjusted R²; the residual fraction is shown outside the circles. Negative fractions are not shown.
